# A Universal Deep Neural Network for In-Depth Cleaning of Single-Cell RNA-Seq Data

**DOI:** 10.1101/2020.12.04.412247

**Authors:** Hui Li, Cory R. Brouwer, Weijun Luo

## Abstract

Single cell RNA sequencing (scRNA-Seq) has been widely used in biomedical research and generated enormous volume and diversity of data. The raw data contain multiple types of noise and technical artifacts and need thorough cleaning. The existing denoising and imputation methods largely focus on a single type of noise (i.e. dropouts) and have strong distribution assumptions which greatly limit their performance and application. We designed and developed the AutoClass model, integrating two deep neural network components, an autoencoder and a classifier, as to maximize both noise removal and signal retention. AutoClass is free of distribution assumptions, hence can effectively clean a wide range of noises and artifacts. AutoClass outperforms the state-of-art methods in multiple types of scRNA-Seq data analyses, including data recovery, differential expression analysis, clustering analysis and batch effect removal. Importantly, AutoClass is robust on key hyperparameter settings including bottleneck layer size, pre-clustering number and classifier weight. We have made AutoClass open source at: https://github.com/datapplab/AutoClass.

## Introduction

scRNA-Seq has been widely adopted in biological and medical research^1-5^ as an ultra-high resolution and ultra-high throughput transcriptome profiling technology. Enormous amount of data has been generated providing great opportunities and challenges in data analytics.

First of all, scRNA-Seq data come with multiple types of noise and quality issues. Some are issues associated with gene expression profiling in general, including RNA amplification bias, uneven library size, sequencing and mapping error, etc. Others are specific to single cell assays. For example, extremely small sample quantity and low RNA capture rate result in large number of false zero expression or dropout^6^. Individual cells vary in differentiation or cell cycle stages^7^, health conditions or stochastic transcription activities, which are biological differences but irrelevant in most studies. In addition, substantial batch effects are frequently observed^8^ due to inconsistence in sample batches and experiments. Most of these noises and variances are not dropout and may follow Gaussian, Poisson or more complex distributions depending on the source of the variances. All of these variances need to be corrected and cleaned so that biologically relevant differences can be reconstructed and analyzed accurately.

Multiple statistical methods have been developed to impute and denoise scRNA-Seq data. Most of these methods rely on distribution assumptions on scRNA-Seq data matrix. For example, deep count autoencoder (DCA)^9^ assumes negative binomial distribution with or without zero inflation, SAVER^10^ assumes negative binomial distribution, and scImpute^11^ uses a mixture of Gaussian and Gamma model. Currently, there is no consensus on the distribution of scRNA-Seq data. Method with inaccurate distribution assumptions^12^ may not denoise properly, but rather introduce new complexities and artifacts. Importantly, these methods largely focus on dropouts and ignore other types of noise and variances, which hinders accurate analysis and interpretation of the data.

To address these issues, we developed AutoClass, a neural network-based method. AutoClass integrates two neural network components: an autoencoder and a classifier (Figure 1a and Methods). The autoencoder itself consists of two parts: an encoder and a decoder. The encoder reduces data dimension and compresses the input data by decreasing hidden layer size (number of neurons). The decoder, in the opposite, expands data dimension and reconstructs the original input data from the compressed data by increasing hidden layer size. Note the encoder and decoder are symmetric in both architecture and function. The data is most compressed at the so-called bottleneck layer between the encoder and the decoder. The autoencoder itself, as an unsupervised data reduction method, is not sufficient in separating signal from noise (Figure 1b). To ensure the encoding process filter out noise and retain signal, we add a classifier branch from the bottleneck layer (Figure 1a and Methods). When cell classes are unknown, virtual class labels are generated by pre-clustering. Therefore, AutoClass is a composite deep neural network with both unsupervised (autoencoder) and supervised (classifier) learning components. AutoClass does not presume any type of data distribution, hence has the potential to correct a wide range noises and non-signal variances. In addition, it can model non-linear relationships between genes with non-linear activation functions. In this study, we extensively evaluated AutoClass against existing methods using multiple simulated and real datasets. We demonstrated AutoClass can better reconstruct scRNA-Seq data and enhance downstream analysis in multiple aspects. In addition, AutoClass is robust over hyperparameter settings and the default setting applies well in various datasets and conditions.

**Figure 1.**
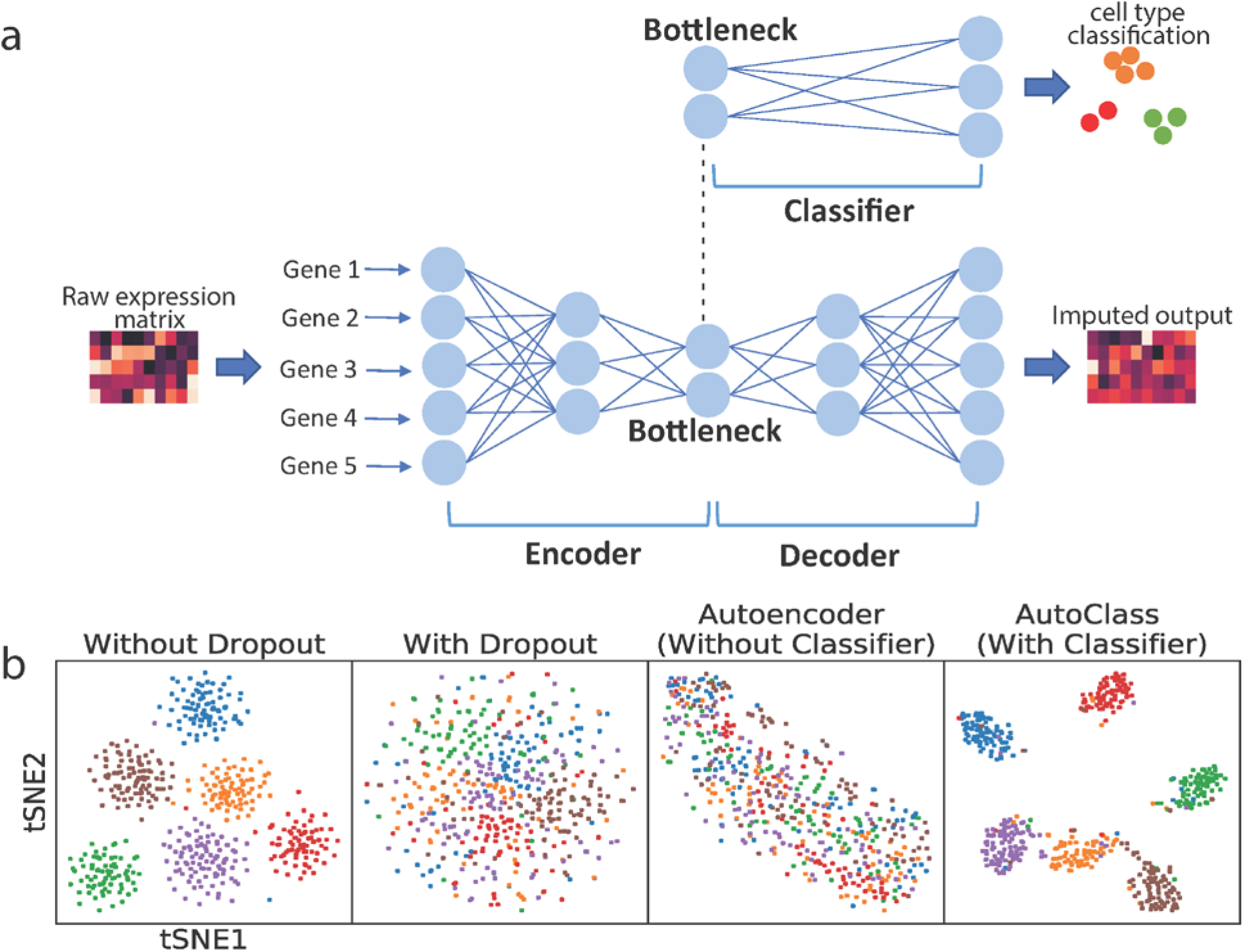
AutoClass integrates a classifier to a regular autoencoder, as to fully reconstruct scRNA-Seq data. **a** AutoClass consists a regular autoencoder and a classifier branch from the bottleneck layer. The input raw expression data is compressed in the encoder, and reconstructed in the decoder, the classifier branch helps to retain signal in data compression. The output of the autoencoder is the desired imputed data (see Methods for details). **b** t-SNE plots of Dataset 1 without dropout, with dropout, with dropout imputed by a regular autoencoder and AutoClass.

## Results

### Validation of the classifier component

The unique part of AutoClass is the classifier branch from the bottleneck layer. Since encoding process losses information in the input data, the classifier branch is added to make sure relevant information or signal is sufficiently retained. To show that the classifier is needed, we simulated a scRNA-Seq Dataset 1 (see Methods and Supplementary Table 2) using Splatter^13^ with 1,000 genes and 500 cells in 6 groups, with and without dropout. Applied both AutoClass and a regular autoencoder without the classifier on the data with dropout, the results are illustrated in two-dimensional t-SNE (see Methods) plots in Figure 1b. AutoClass but not the regular autoencoder was able to recover cell type pattern, indicating the classifier component is necessary for reconstructing scRNA-Seq data.

### Gene expression data recovery

We evaluated expression value recovery on simulated scRNA-Seq data with different noise types or distributions. We generated and scRNA-Seq dataset using Splatter with 500 cells, 1000 genes in five cell groups with (raw data, Dataset 2) and without dropout (true data). From the same true data, we also generated 5 additional raw datasets by adding noise following different distributions which are representative and commonly seen, including random uniform (Dataset 3), Gaussian (Dataset 4), Gamma (Dataset 5), Poisson (Dataset 6) and negative binomial (Dataset 7) (details in Methods and Supplement Table 2 and 3).

As expected, dropout noise greatly reduced the data quality and obscured the signal or biological differences such as distinction between cell types (Figure 2a). All other noise types had similar effect on the data (Figure2b-2c and Supplementary Figure 1). With t-SNE transformation on Dataset 2-7, the true data without noise showed distinct cell types, but not the raw data with noises (Figure 2a-c and Supplementary Figure 1). The average Silhouette width^14^ (ASW) on the t-SNE plot is a measurement of distance between groups, ranges from −1 to 1, where higher values indicate more confident clustering. ASW dropped greatly from 0.64 to around 0 in all raw datasets. After imputation by AutoClass, the cell type pattern was recovered and ASW increased back substantially to 0.2-0.5. In contrast, all published control methods (DCA, MAGIC^15^, scImpute and SAVER) were unable to recover the original cell type pattern (Figure 2a-c and Supplementary Figure 1) and ASW scores remained low (Figure 2d) for all noise types.

**Figure 2.**
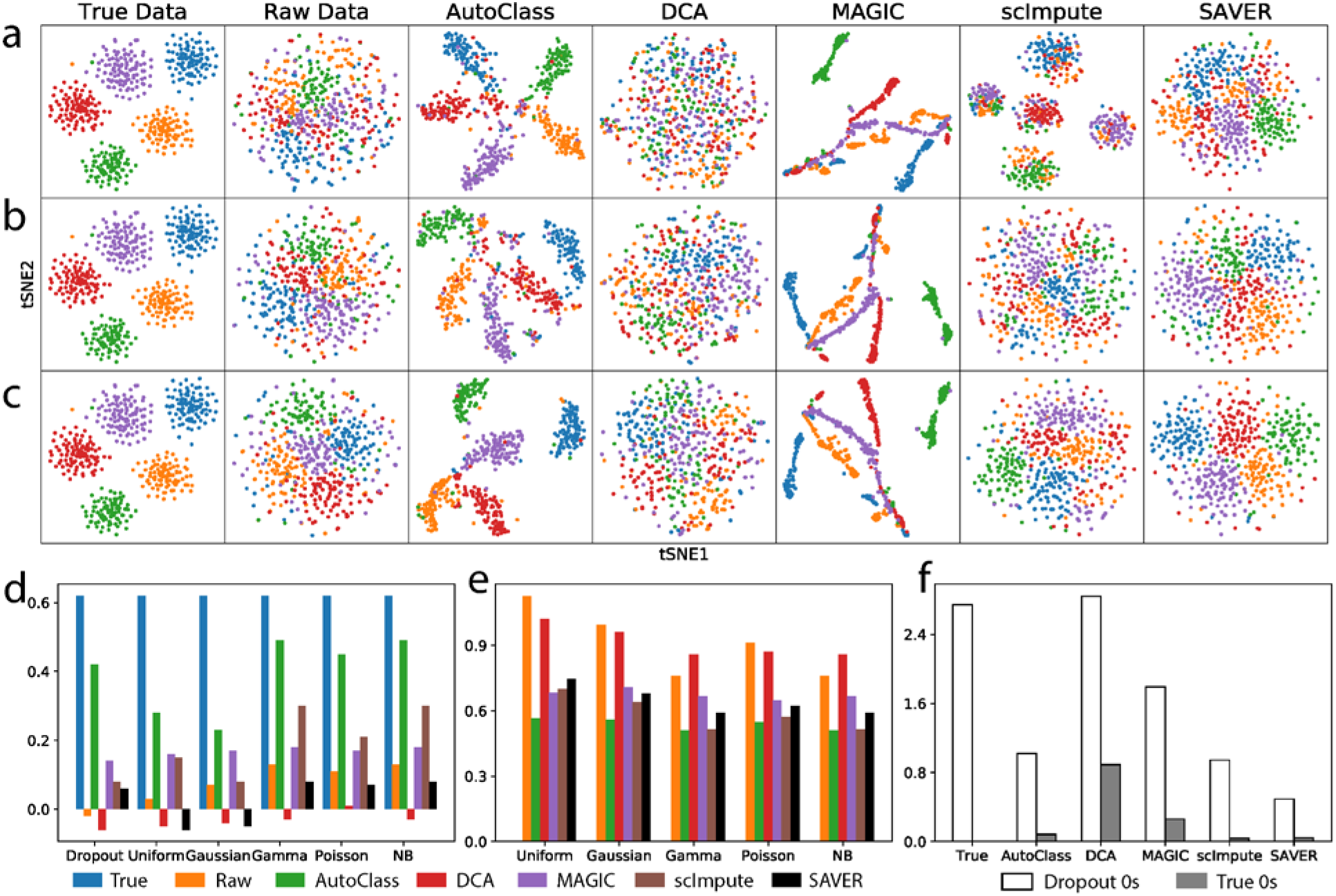
Gene expression data recovery after imputation. **a**, **b** and **c** t-SNE plots for Dataset 2 (dropout noise), Dataset 4 (Gaussian noise) and Dataset 7 (negative binomial noise), respectively. **d** Average Silhouette width based on t-SNE plot for Dataset 2-7. **e** Mean squared error between true data and imputed data for Dataset 3-7. **f** Average recovered values of dropout 0s and true 0s for different imputation methods.

We also measure the data recovery quality using other metrics. The mean squared error (MSE) between the true data and imputed/denoised data for Dataset 3-7 (5 noise types other than dropout) were also computed (Figure 2e). Among the 5 tested methods, AutoClass consistently achieved the smallest MSE for all noise types (Figure 2e). Dropout noise (Dataset 2) is very different from all other noise types (Dataset 3-7) in both distribution form and generation mechanism, and MSE was not an informative measurement of data recovery. We computed the average recovered values of dropout zeros and those of true zeros (Figure 2f) instead. An ideal imputation method can distinguish between these two types of zeros, i.e. impute dropout zeros while retain true zeros (Figure 2f). While SAVER was too conservative in imputing both types of 0 values, DCA and MAGIC were too aggressive. AutoClass and scImpute both achieved good balance between imputing dropout 0s and retaining true 0s, yet only the former but bot the later was able to recover the biological difference or distinct cell type clustering (Figure 2a-c and Supplementary Figure1).

### Differential expression analysis

Differential expression (DE) analysis is by far the most common analysis of scRNA-Seq and gene expression data. To study the performance of AutoClass in DE analysis, we simulated a scRNA-Seq Dataset 8 using Splatter with 1,000 genes and 500 cells in two cell groups. Here the ground truth of truly differentially expressed genes is known. We applied Two-sample T-test to the true, raw and imputed data using different methods. The median value of t-statistics for the truly differentially expressed genes dropped from 5.79 in the true data to 2.11 in the raw data, and increased back to 5.86 upon imputation by AutoClass, which was almost the same as in the true data and higher than in all control methods (Figure 3 a - b). As shown by ROC curves and area under the curves (AUC), AutoClass also was the best at balancing true positives and false negatives (Figure 3 c - d).

**Figure 3.**
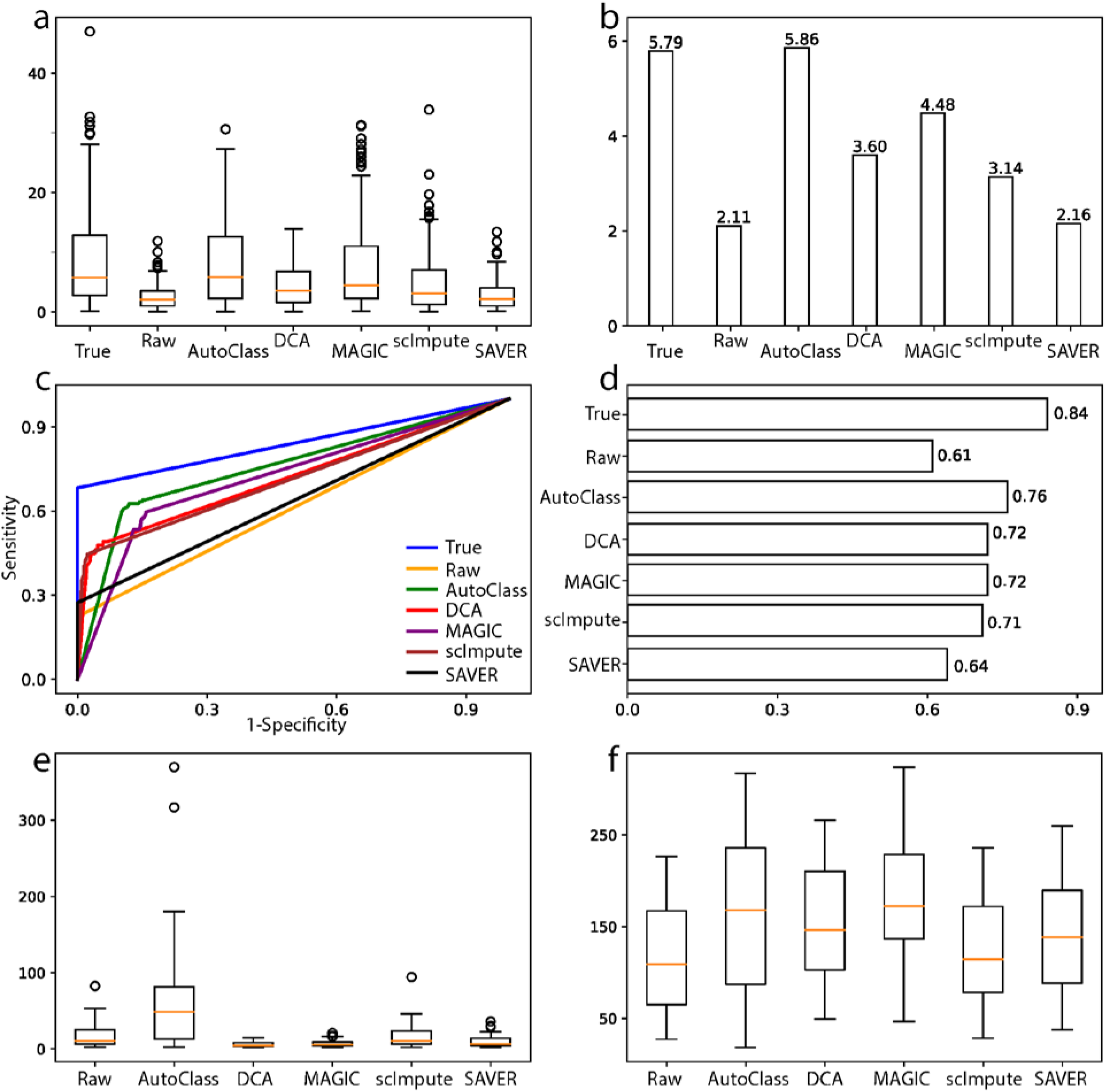
Differential expression analysis and marker gene analysis. **a** and **b** T-statistics and their median values for truly differentially expressed genes in Dataset 8. **c** and **d** ROC curves and areas under the ROC curves for Dataset 8. **e** and **f** Fold changes and T-statistics of marker genes in the Baron dataset.

Similarly, AutoClass can improve DE analysis in data with Gaussian noise. We manually added Gaussian noise to the true data of Dataset 8 to generate the raw data of Dataset 9. The DE analysis results can be found in Supplementary Figure 2.

AutoClass also improves marker gene expression analysis. Baron dataset^16^ provides known marker gene lists for related cell types in pancreatic islets. AutoClass imputed data increased both t-statistics and fold changes of the marker genes (Figure 3 e - f).

### Clustering analysis

Clustering analysis is frequently done on scRNA-seq data as to identify cell types or subpopulations. To evaluate AutoClass for clustering analysis, we used four real datasets, including two small datasets: the Buettner dataset^2^ (182 cells) and the Usoskin dataset^17^ (622 cells) and two large datasets: the Lake dataset^18^ (8,592 cells) and the Zeisel dataset^19^ (3,005 cells). Detailed information for these datasets can be found in Methods and Supplementary Table 1.

We compared K-means clustering results on the 200 highest variable genes. The ground truth or the actual number of cell types were used as number of clusters. Clustering results were evaluated by four different metrics: adjusted Rand index^20^ (ARI), Jaccard Index^21^ (JI), normalized mutual information^22^ (NMI) and purity score^23^ (PS). All of them range from 0 to 1, with 1 indicating a perfect match to the true groups. AutoClass is the only method improving all four metrics from the raw data. In addition, AutoClass achieved the best clustering results for 3 out of 4 datasets (Table 1).

**Table 1.** Evaluation of clustering results of four real scRNA-Seq datasets. The four metrics are adjusted Rand index (ARI), Jaccard Index (JI), normalized mutual information (NMI) and purity score (PS). Highest value in each row was highlighted in boldface.

| metric | dataset | Raw | AutoClass | DCA | MAGIC | scImpute | SAVER |
| --- | --- | --- | --- | --- | --- | --- | --- |
| ARI | Buettner | 0.023 | **0.372** | 0.288 | 0.213 | 0.039 | 0.016 |
|  | Usoskin | 0.221 | **0.869** | 0.234 | 0.813 | 0.067 | 0.317 |
|  | Lake | 0.403 | 0.557 | **0.572** | 0.440 | 0.313 | 0.465 |
|  | Zeisel | 0.737 | **0.793** | 0.753 | 0.433 | 0.623 | 0.763 |
| JI | Buettner | 0.242 | **0.409** | 0.363 | 0.368 | 0.262 | 0.247 |
|  | Usoskin | 0.324 | **0.830** | 0.284 | 0.764 | 0.266 | 0.351 |
|  | Lake | 0.323 | 0.439 | **0.453** | 0.346 | 0.254 | 0.364 |
|  | Zeisel | 0.646 | **0.713** | 0.664 | 0.370 | 0.679 | 0.677 |
| NMI | Buettner | 0.035 | **0.395** | 0.333 | 0.335 | 0.075 | 0.038 |
|  | Usoskin | 0.225 | **0.829** | 0.253 | 0.771 | 0.048 | 0.431 |
|  | Lake | 0.611 | 0.667 | **0.676** | 0.601 | 0.500 | 0.642 |
|  | Zeisel | 0.747 | **0.784** | 0.746 | 0.598 | 0.798 | 0.762 |
| PS | Buettner | 0.434 | **0.720** | 0.648 | 0.599 | 0.445 | 0.423 |
|  | Usoskin | 0.545 | **0.937** | 0.579 | 0.913 | 0.416 | 0.682 |
|  | Lake | 0.723 | **0.772** | 0.766 | 0.693 | 0.610 | 0.742 |
|  | Zeisel | 0.894 | **0.917** | 0.880 | 0.763 | 0.548 | 0.897 |

For the Usoskin dataset, out of all tested methods, only AutoClass and MAGIC reconstructed distinct clusters (Figure 4a). But MAGIC likely generated false positive signals, given that the between-group cell-to-cell correlation are almost the same as within-group correlation, and both are close to 1 (Figure 4b). AutoClass was the only method differentiating within-group vs between-group correlation as informative metrics for signal vs noise (Figure 4b).

**Figure 4.**
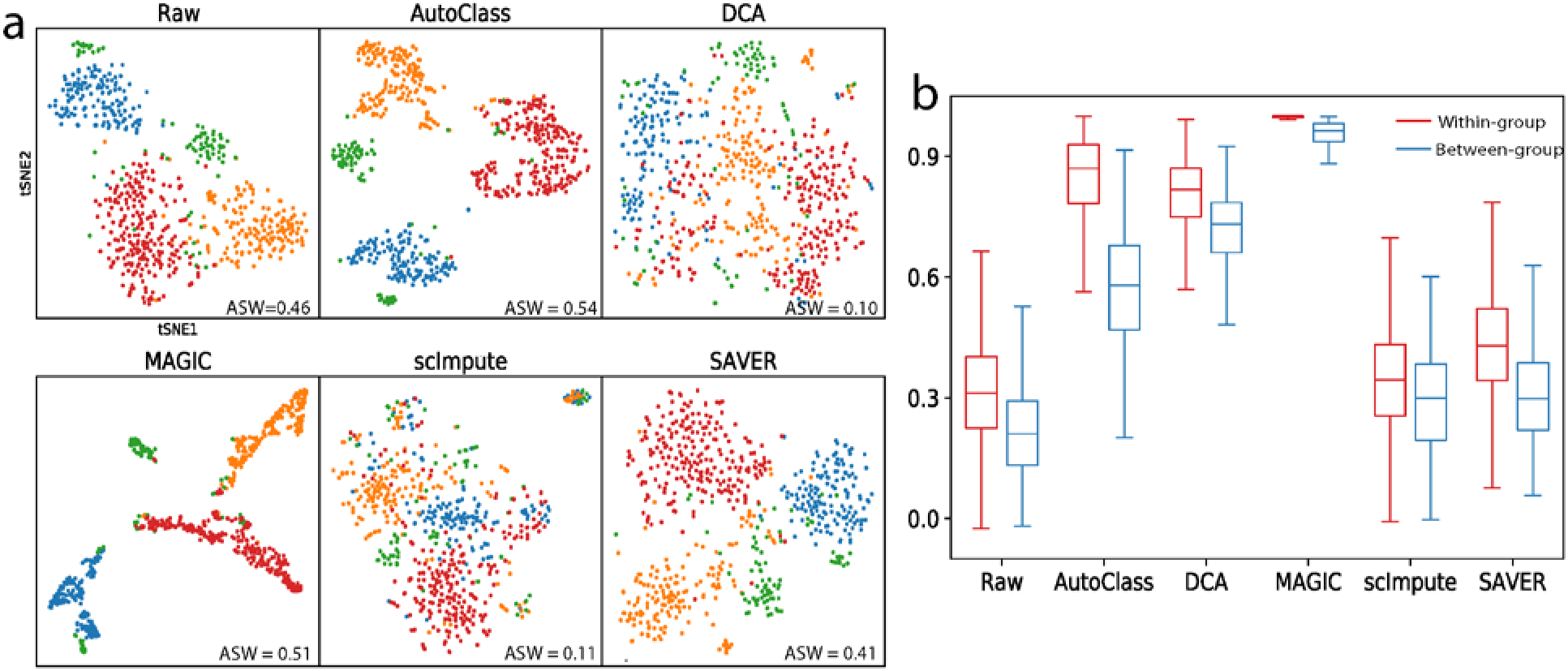
Imputation results for Usoskin data. **a** t-SNE plots for raw and imputed data. **b** Within-group and between-group cell-to-cell correlation for raw and imputed data.

### Batch effect removal

Batch effect rises from different individual cell donors, sample groups, or experiment conditions and can severely affect downstream analysis. We analyzed two real datasets with major batch effect. The Villani dataset^24^ sequenced 768 human blood dendritic cells (DC) in 2 batched using Smart-Seq2. The Baron dataset includes 7,162 pancreatic islet cells from 3 healthy individuals.

Similar to Tran et al.^8^, we evaluated the performance of batch effect correction as the ability to merge different batches of the same cell type while keeping different cell types separate. We did t-SNE transformation on the data first (Figure 5a and Supplementary Figure 3), then applied the four metrics above mentioned, i.e. ASW, ARI, NMI and PS on both cell types and batches. While cell-type-level metrics measure cell type separation, 1 - batch level metrics measure the merging between batches of same cell type (Figure 5b and Supplementary Figure 4).

**Figure 5.**
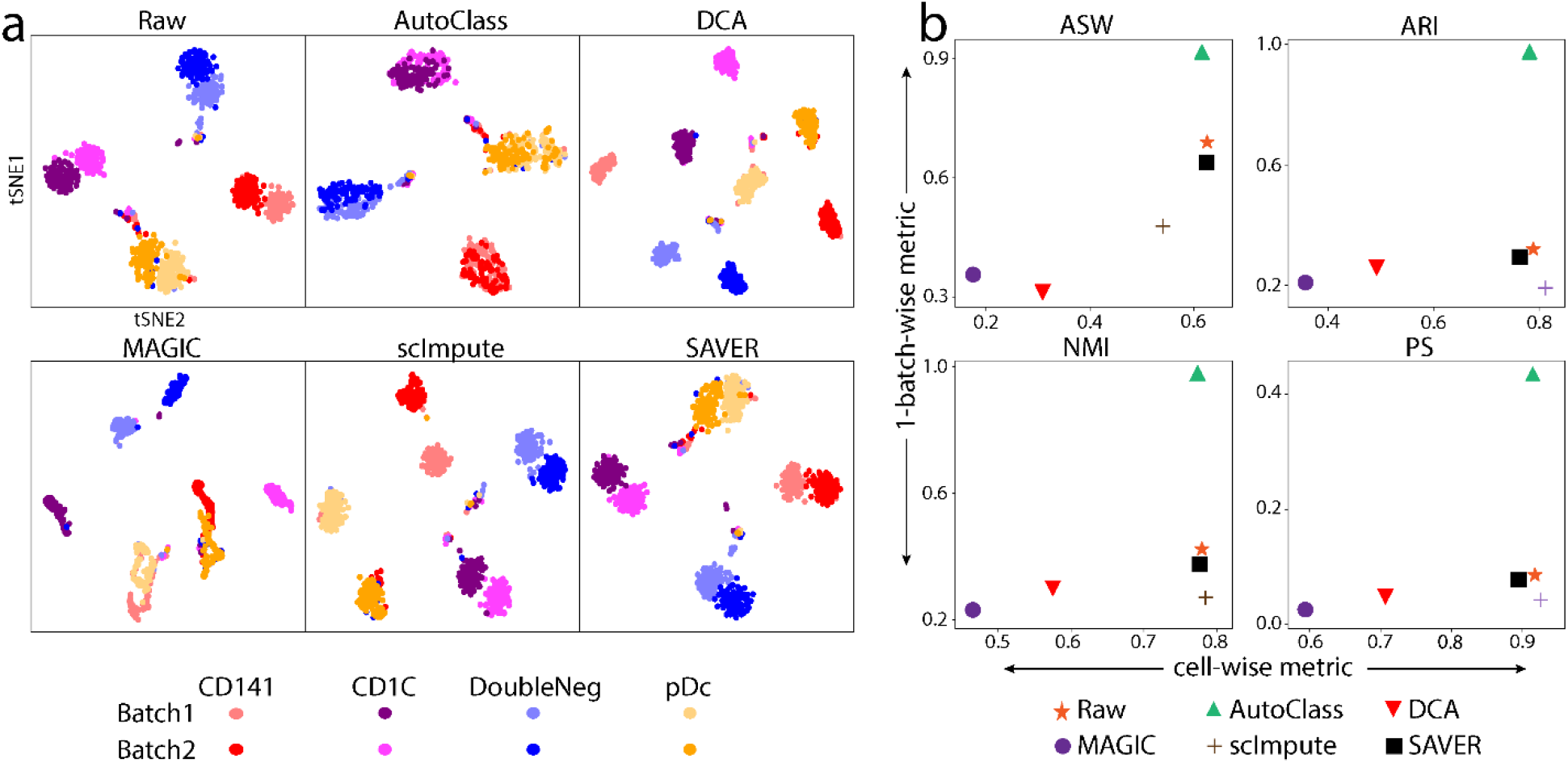
Batch effect in Villani dataset. **a** t-SNE plots for raw and imputed data. **b** Evaluation of batch and cell type separation in raw and imputed data by four different metrics: average Silhouette width, adjusted Rand index, normalized mutual information and purity score. Good performance is show as big values in both X-(cell type separation) and Y-(batch effect removal) axes.

In the Villani dataset (Figure 5), the raw data shows clear separation in both cell types and sample batches. After imputation by AutoClass, while cell types remained well separated, the two batches were evenly mixed up within each cell type. In contrast, SAVER failed to reduce the batch effect, while all other methods even aggravated it (Figure 5).

Note that AutoClass corrects the batch effect without knowing the actual number of cell types. Here, we used the default number of clusters in the pre-clustering step, i.e., [8, 9, 10] (see Methods). This is close to the number of spurious groups counting batches (i.e., 8), but far away from the actual number of cell types, or 4. In other words, AutoClass was not misled by the pre-clustering number and correctly recovered the actual cluster number.

In Baron dataset (Supplementary Figure 3 and 4), AutoClass reduced the batch effect and increased cell type separation simultaneously with the default pre-clustering number too. MAGIC dramatically reduced the differences in both batches and cell type. The batch effect correction by other methods were limited.

### Robustness over major hyperparameters

AutoClass, as a composite deep neural network, has multiple hyperparameters. Among them, the most important ones are bottleneck layer size, number of pre-clusters and classifier weight. Bottleneck layer plays an important role in autoencoders, it is the narrowest part of the network and the size (number of neurons) controls how much the input data is compressed. The number of clusters (*K*) in the pre-clustering step is specific to the classifier of AutoClass. AutoClass uses three consecutive cluster numbers [*K* — 1, *K, K* + 1], and the final imputation output is the average over three predictions using these three clustering numbers (see Methods). In addition, the classifier weight *w* (see Equation (4) and Methods) is another AutoClass specific hyperparameter which balance the ratio between autoencoder loss and the classifier loss.

AutoClass is robust over a wide range of bottleneck layer sizes, pre-clustering *K* values (Figure 6 and Supplementary Figure 5-6) and classifier weight *w* (Supplementary Figure 7). The t-SNE clustering patterns, clustering metrics (ASW and ARI), MSE and imputed dropout0s/true 0s ratio remained the same when bottleneck layer size increase from 16 to 256 (Figure 6a, 6c, Supplementary Figure 6a). However, these results or metrics varied heavily in the same analysis using DCA, another autoencoder based method (Figure 6b, 6c, Supplementary Figure 6a). Likewise, AutoClass also achieved stable results over the range of *K* values - 4-8 (Figure 6d, Supplementary Figure 5, Supplementary Figure 6b) and the range of classifier weight *w* values - 0.1-0.9 (Supplementary Figure 7).

**Figure 6.**
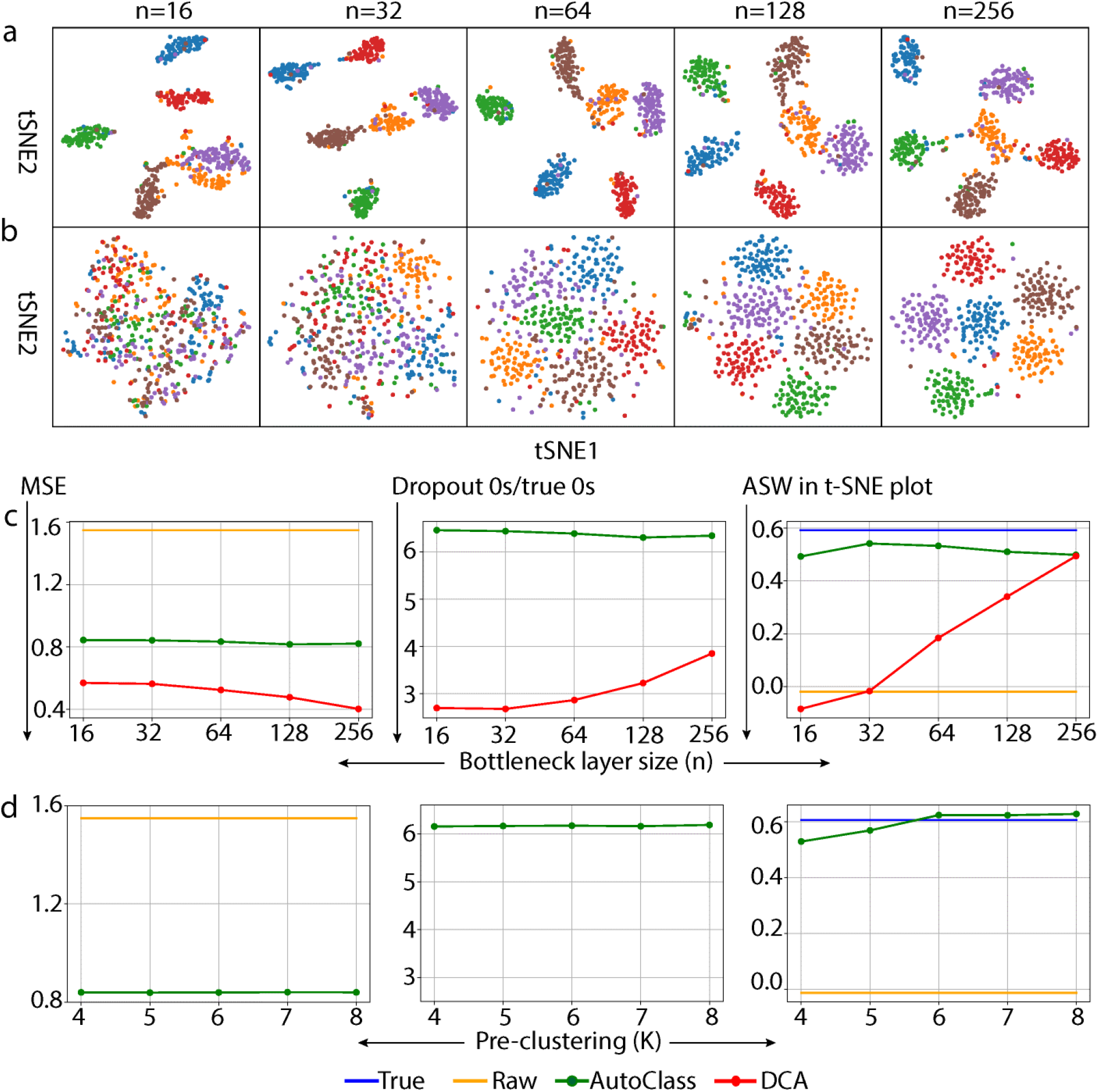
Impact of bottleneck layer size and the number of clusters in the pre-clustering in Dataset 1. **a** and **b** t-SNE plots of data imputed by AutoClass (**a**) and DCA (**b**) with different bottleneck sizes (n). **c** and **d** Imputation results by AutoClass with bottleneck layer size (**c**) and the number of clusters in the pre-clustering (**d**).

## Discussion

In this work, we proposed and developed a deep learning-based method AutoClass for thorough cleaning of scRNA-Seq data. AutoClass integrates two neural network components, an autoencoder and a classifier. This composite network architecture is essential for filtering out noise and retaining signal effectively. Unlike many other scRNA-Seq imputation methods, AutoClass does not rely on any distribution assumption, and fully counts the non-linear interactions between genes. With these properties, AutoClass effectively models and cleans a wide range of noises and artifacts in scRNA-Seq data including dropouts, random uniform, Gaussian, Gamma, Poisson and negative binomial noises, as well as batch effects. These are the most common and representative types of noises and artifacts. Any other types not directly tested would likely be cleaned with the same efficiency because they are similar in distribution and source and AutoClass has no assumption on the noise forms. Such in-depth cleaning led to consistent and substantial improvement of the data quality and downstream analyses including differential expression and clustering, as shown by a range of experiments with both simulated and real datasets.

Hyperparameter tuning is an important yet tedious step for training neural network models. Inadequate tuning of hyperparameters may lead to suboptimal results. Remarkably, AutoClass is robust with key hyperparameters including bottleneck layer size (*n*), pre-clustering number (*K*) and classifier weight (*w*). The default setting with *n* = 128, *K* =9, *w* = 0.9 works well for most scRNA-Seq datasets and conditions. This robustness makes AutoClass an appealing method for both performance and practical uses.

## Methods

### Architecture of AutoClass

AutoClass integrates two neural network components, an autoencoder and a classifier, to impute scRNA-seq data (Figure 1a). The classifier branch is necessary to preserve signals or biological differences (cell type patterns etc.) from loss in data compression by the encoder.

When cell classes are unknown, virtual class labels are generated by pre-clustering using *K*-means method. The total loss of the entire network is the weighted sum of classifier loss (cross-entropy or CE) and the autoencoder loss (MSE). The activation functions in the hidden layers are all rectified linear unit (ReLU), the activation functions for the output layer of autoencoder and classifier are SoftPlus and SoftMax, respectively.

The formulation of AutoClass architecture is:

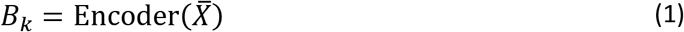

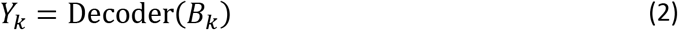

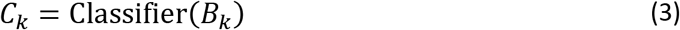

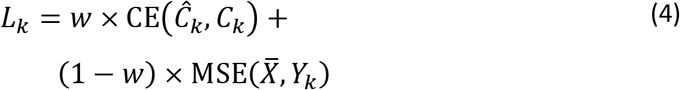

Where *B_k_, Y_k_, C_k_* and *L_k_* are the bottleneck representation, the output of the decoder hence the autoencoder, the output of the classifier and total loss, respectively. 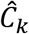 is the pre-clustering cell type labels for *k* clusters. 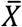 is the input of AutoClass, and has been normalized over library size and followed by a log_2_ transformation with pseudo count 1:

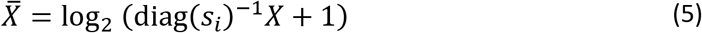

*X* is the raw count matrix and the size factor *s_i_* for cell *i* is equal to the library size divided by the median library size across cells. Library size is defined as the total number of counts per cell.

The final imputed data is the average prediction of the autoencoder over different cluster numbers in pre-clustering:

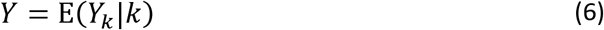

For all datasets in this manuscript, we used 3 consecutive cluster numbers, or *k* = [*K* — 1, *K, K* + 1], the default value is *K* = 9. The final imputation result was the average results over different *K_s_*.

### AutoClass Implementation and hyperparameter settings

AutoClass is implemented in Python 3 with Keras. *Adam* is used for optimizer with default learning rate 0.001. Learning rate is multiplied by 0.1 if validation loss does not improve for 15 epochs. The training stops if there is no improvement for 30 epochs.

Although AutoClass works well for small bottleneck layer sizes (*n* = 16, 32 or similar), we set the default value to be *n* = 128, as be conservative and to avoid potential information loss in data compression. This default value was used in all datasets in this paper.

AutoClass is stable over different choices of *K* in pre-clustering as long as *K* is not extremely far away from the true number of cell clusters. The default value *K* =9 was used in all datasets in this paper except simulated Dataset 8 and 9, since the true number of cell clusters in these two datasets is 2 which is far smaller than default value 9. Hyperparameter *K* can be chosen based on prior knowledge of the data or statistical methods like elbow method^25^ and Silhouette method^14^. *K* used in Dataset 8 and 9 was the average of estimations by elbow method and Silhouette method.

AutoClass is stable on classifier weight *w* in the range of 0.1-0.9 (supplementary Figure 7). We found that in general classification loss is far smaller than reconstruction loss (Supplementary Figure 8), to have a better balance between those two losses, we set the default value to be *w* = 0.9. This default value was used in all the datasets in this paper.

In addition, overfitting is a common problem in neural network models. Dropout of neurons^26^ in the bottleneck layer is used in AutoClass to prevent overfitting. Interestingly, a relatively high dropout rate in AutoClass also helps to correct batch effect. In the batch effect removal analyses, we set dropout rate to be 0.5 in AutoClass, and to be fair, also in DCA. But DCA was unable to remove batch effect (Figure 5, Supplementary Figure 3-4). The default dropout rate 0.1 in AutoClass was used in all the other datasets and analyses in this paper.

AutoClass hyperparameter settings for all the datasets can be found in Supplementary Table 1-3.

### Analysis details

#### Noise types other than dropout

Dataset 3-7 and Dataset 9 were generated by manually adding noise to the true data of Dataset 2 and Dataset 8, respectively. The noise was first generated by *Python numpy.random* package with different noise distributions (details in Supplementary Table 3), and then centered (so that noise mean is 0). The noise was then added to true data, all values were rounded to be integers and negative values set to 0, since scRNA-Seq data raw counts are positive integers.

#### Highly variable genes

The highly variable genes in each dataset are ranked by the ratio between gene-wise variance vs mean computed from non-zero values.

#### t-Distributed stochastic neighbor embedding (t-SNE)

We applied t-SNE^27^ to visualize datasets. We first reduce the number of data dimensions by using the top 50 principle components, and then use *TSNE* function in the *sklearn.manifold* package with default settings to further reduce the dimension to 2 for visualization.

#### Batch effect removal score

Four clustering metrics ASW, ARI, NMI and PS were used to measure the performance of batch effect correction. We applied ASW to the t-SNE transformed data, and batch effect removal was scored by both cell-type-wise ASW vs 1 – batch-wise ASW (Figure 5b and Supplementary Figure 4). Higher values in both dimensions together denote better batch effect removal. ARI, NMI and PS metrics are used and plotted in the same fashion as ASW. To compute ARI, NMI and PS, K-means clustering was performed first to obtain cluster labels, which were then compared to batch labels and cell type labels. The batch indices were computed for each individual cell type first, and take weighted sum across cell types. The weight for each cell type is proportional to the number of cells.

### Control methods

DCA (version 0.2) was downloaded from https://github.com/theislab/dca

Magic (version 0.1.0) was downloaded from https://github.com/KrishnaswamyLab/MAGIC

scImpute (version 0.0.5) was downloaded from https://github.com/Vivianstats/scImpute.

SAVER (version 0.3.0) was downloaded from https://github.com/mohuangx/SAVER.

### Real scRNA-seq datasets

We collected and analyzed multiple real scRNA datasets from published studies. These datasets have been well established, widely used and tested as shown in literature. While major technical attributes are summarized in Supplementary Table 1, below are more details.

#### Baron Study

Human pancreatic islets cells data were obtained from 3 healthy individuals, which provided gene expression profiles for 17,434 genes in 7,729 cells. We filtered out genes expressed in less than 5 cells, removed cell types less than 1% of the cell population. Analysis was restricted to top 1,000 highly variable genes. Final dataset contained 7,162 cells with 8 different cell types.

The raw counts data are available at https://shenorrlab.github.io/bseqsc/vignettes/pages/data.html.

#### Villani Study

The human blood dendritic data contained 26,593 genes in 1,140 cells. We kept batch 1 (plate id: P10, P7, P8 and P9) batch 2 (plate id P3, P4, P13, P14) cells, and filtered out genes expressed in less than 5 cells. Analysis was restricted to top 1,000 highly variable genes. Final dataset contained 768 cells with 4 different cell types in 2 batches. The raw data are available at GEO accession GSE80171.

#### Lake Study

Human brain frontal cortex data contained 34,305 genes in 10,319 cells. We filtered out genes expressed in less than 5 cells, removed cell types h less than 3% of the cell population. Analysis was restricted to top 1,000 highly variable genes. Final dataset contained 8,592 cells with 11 different cell types. The raw data are available at GEO accession GSE97930.

#### Zeisel Study

Mouse cortex and hippocampus data contained 19,972 genes in 3,005 cells. We filtered out genes expressed in less than 5 cells. Analysis was restricted to top 1,000 highly variable genes. Final dataset contained 3,005 cells with 9 different cell types. Annotated data are available at http://linnarssonlab.org/cortex.

#### Buettner Study

Mouse embryonic stem cells contained 8,989 genes in 182 cells. We filtered out genes expressed in less than 5 cells. Final dataset contained 8,985 genes and 182 cells in 3 cells lines. The full dataset was deposited at ArrayExpress: E-MTAB-2805. The normalized data can be obtained from https://www.nature.com/articles/nbt.3102.

#### Usoskin Study

Neuronal data contained 17,772 genes in 622 cells. We filtered out genes expressed in less than 5 cells. Analysis was restricted to top 1,000 highly variable genes. Final dataset contains 622 cells with 4 different cell types. The normalized data can be obtained from https://www.nature.com/articles/nbt.3102.

### Simulated scRNA-seq datasets

Splatter R (version v1.2.2) package was used to simulate scRNA-seq datasets with dropout values. Gaussian noise was manually added when needed. Genes expressed in less than 3 cells were filtered out before analysis. The parameter settings for simulation are summarized in Supplementary Table 2 and 3.

### Code availability

AutoClass python module, documentation, tutorial with example, and code to reproduce the main results in the manuscript are available online: https://github.com/datapplab/AutoClass.

## Data availability

All simulated datasets can be generated using the parameters specified in the Simulated scRNA-Seq datasets subsection, all the real datasets are publicly available as mentioned in the Real scRNA-Seq datasets subsection. In addition, multiple simulated and real datasets were provided in the GitHub repository above as demo datasets, ready for analysis.

## References

1. Griffiths, J.A., Scialdone, A. & Marioni, J.C. Using single-cell genomics to understand developmental processes and cell fate decisions. Mol Syst Biol 14, e8046 (2018).

2. Buettner, F. et al. Computational analysis of cell-to-cell heterogeneity in single-cell RNA-sequencing data reveals hidden subpopulations of cells. Nat Biotechnol 33, 155–160 (2015).

3. Papalexi, E. & Satija, R. Single-cell RNA sequencing to explore immune cell heterogeneity. Nat Rev Immunol 18, 35–45 (2018).

4. Keren-Shaul, H. et al. A unique microglia type associated with restricting development of Alzheimer’s disease. Cell 169, 1276–1290 e1217 (2017).

5. Stubbington, M.J.T., Rozenblatt-Rosen, O., Regev, A. & Teichmann, S.A. Single-cell transcriptomics to explore the immune system in health and disease. Science 358, 58–63 (2017).

6. Hicks, S.C., Townes, F.W., Teng, M. & Irizarry, R.A. Missing data and technical variability in single-cell RNA-sequencing experiments. Biostatistics 19, 562–578 (2018).

7. Trapnell, C. et al. The dynamics and regulators of cell fate decisions are revealed by pseudotemporal ordering of single cells. Nat Biotechnol 32, 381–386 (2014).

8. Tran, H.T.N. et al. A benchmark of batch-effect correction methods for single-cell RNA sequencing data. Genome Biol 21, 12 (2020).

9. Eraslan, G., Simon, L.M., Mircea, M., Mueller, N.S. & Theis, F.J. Single-cell RNA-seq denoising using a deep count autoencoder. Nat Commun 10, 390 (2019).

10. Huang, M. et al. SAVER: gene expression recovery for single-cell RNA sequencing. Nat Methods 15, 539–542 (2018).

11. Li, W.V. & Li, J.J. An accurate and robust imputation method scImpute for single-cell RNA-seq data. Nat Commun 9, 997 (2018).

12. Svensson, V. Droplet scRNA-seq is not zero-inflated. Nat Biotechnol 38, 147–150 (2020).

13. Zappia, L., Phipson, B. & Oshlack, A. Splatter: simulation of single-cell RNA sequencing data. Genome Biol 18, 174 (2017).

14. Rousseeuw, P.J. Silhouettes: a graphical aid to the interpretation and validation of cluster analysis Journal of Computational and Applied Mathematics 20, 53–65 (1987).

15. Dijk, D.v. et al. Recovering gene interactions from single-cell data using data diffusion. Cell 174, 716–729 (2017).

16. Baron, M. et al. A single-cell transcriptomic map of the human and mouse pancreas reveals Inter-and Intra-cell population structure. Cell Syst 3, 346–360 e344 (2016).

17. Usoskin, D. et al. Unbiased classification of sensory neuron types by large-scale single-cell RNA sequencing. Nat Neurosci 18, 145–153 (2015).

18. Lake, B.B. et al. Integrative single-cell analysis of transcriptional and epigenetic states in the human adult brain. Nat Biotechnol 36, 70–80 (2018).

19. Zeisel, A. et al. Brain structure. Cell types in the mouse cortex and hippocampus revealed by single-cell RNA-seq. Science 347, 1138–1142 (2015).

20. Hubert, L. & Arabie, P. Comparing partitions. Journal of classification 2, 193–218 (1985).

21. Leydesdorff, L. On the normalization and visualization of author co-citation data: Salton’s Cosine versus the Jaccard index. Journal of the American Society for Information Science and Technology 59, 77–85 (2007).

22. Estevez, P.A., Tesmer, M., Perez, C.A. & Zurada, J.M. Normalized mutual information feature selection. IEEE Trans Neural Netw 20, 189–201 (2009).

23. Manning, C.D., Raghavan, P. & Schutze, H. Introduction To Information Retrieval. (Cambridge Univ. Press, Cambridge; 2008).

24. Villani, A.C. et al. Single-cell RNA-seq reveals new types of human blood dendritic cells, monocytes, and progenitors. Science 356(2017).

25. Ketchen, D.J. & Shook, C.L. The application of cluster Analysis in strategic management research: an analysis and critique. Strategic Management 17, 441–458 (1996).

26. Srivastava, N., Hinton, G., Krizhevsky, A., Sutskever, I. & Salakhutdinov, R. Dropout: a simple way to prevent neural networks from overfitting. J Mach Learn Res 15(2014).

27. Maaten, L.v.d. & Hinton, G. Visualizing data using t-SNE. Journal of machine learning research 9, 2579–2605 (2008).

