## Supplementary Figures & Tables for "A Universal Deep Neural Network for In-Depth Cleaning of Single-Cell RNA-Seq Data"

### Supplementary Materials

Supplementary Figures 1 to 8

Supplementary Tables 1 to 3

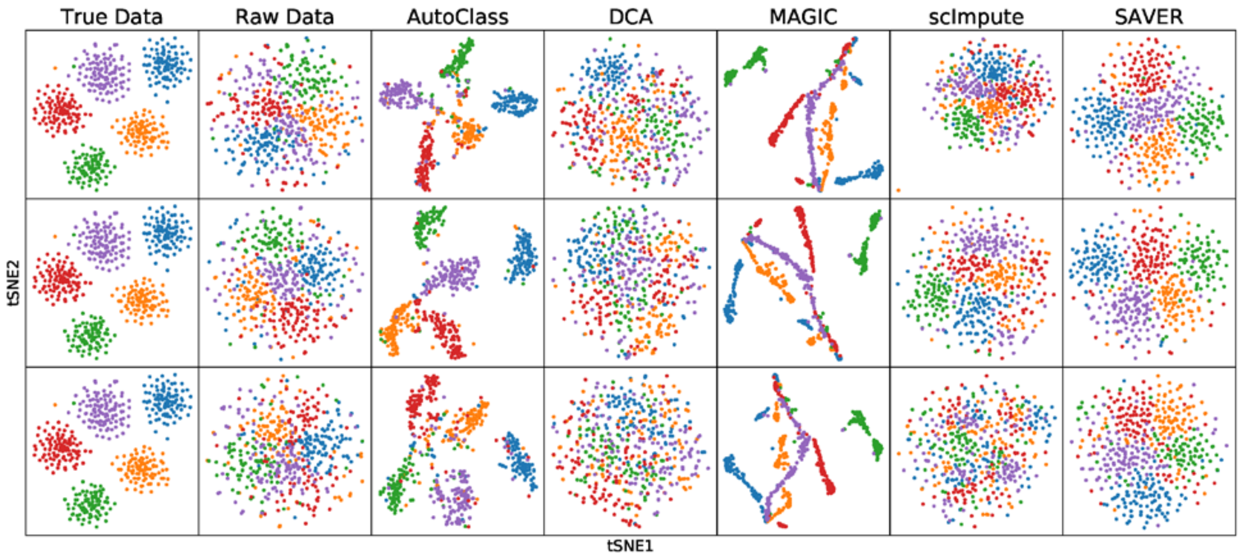


**Supplementary Figure 1** t-SNE plots for Dataset 3 (first row, random uniform noise), Dataset 5 (second row, Gamma noise) and Dataset 6 (third row, Poisson noise). Experiments settings are similar to those in Figure 2.

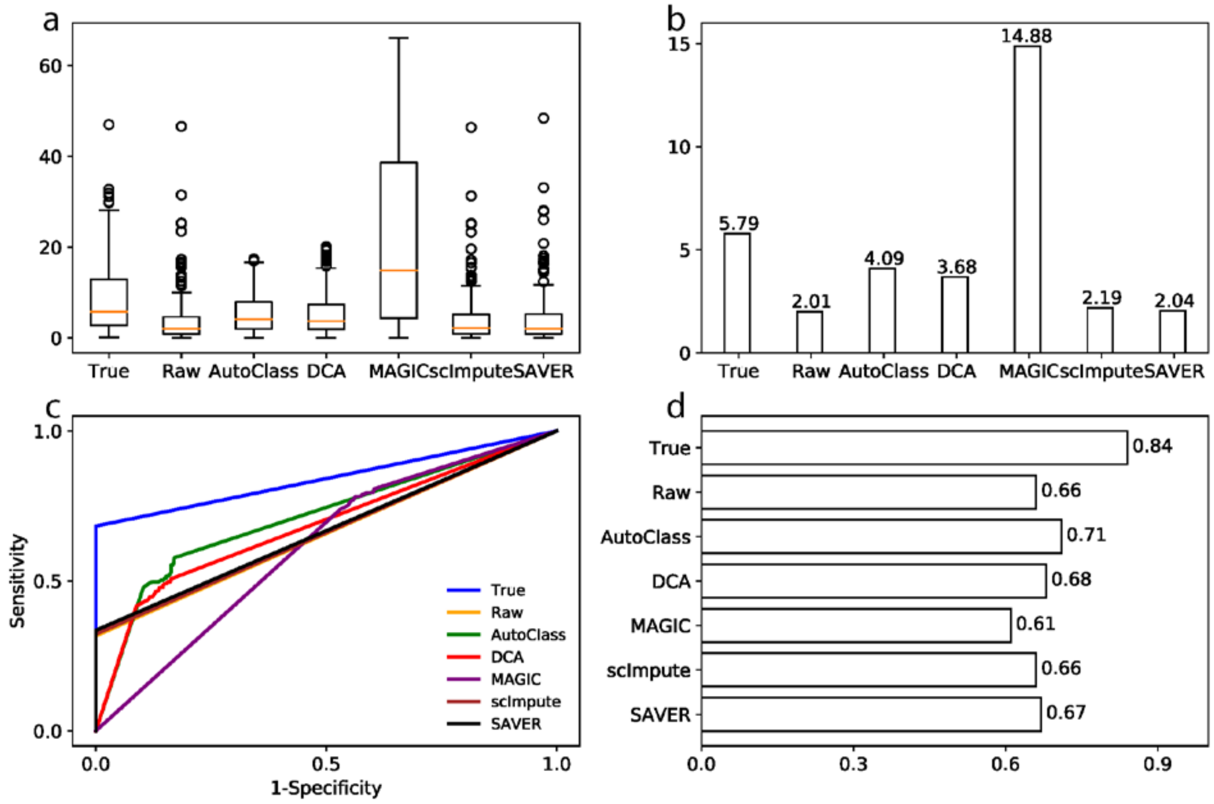


**Supplementary Figure 2** Differential expression analysis for Dataset 9 (Gaussian noise). **a** and **b** T-statistics and their median for truly differentially expressed genes. **c** and **d** ROC curves and areas under the ROC curves. Experiments settings are similar to those in Figure 3 a-d.

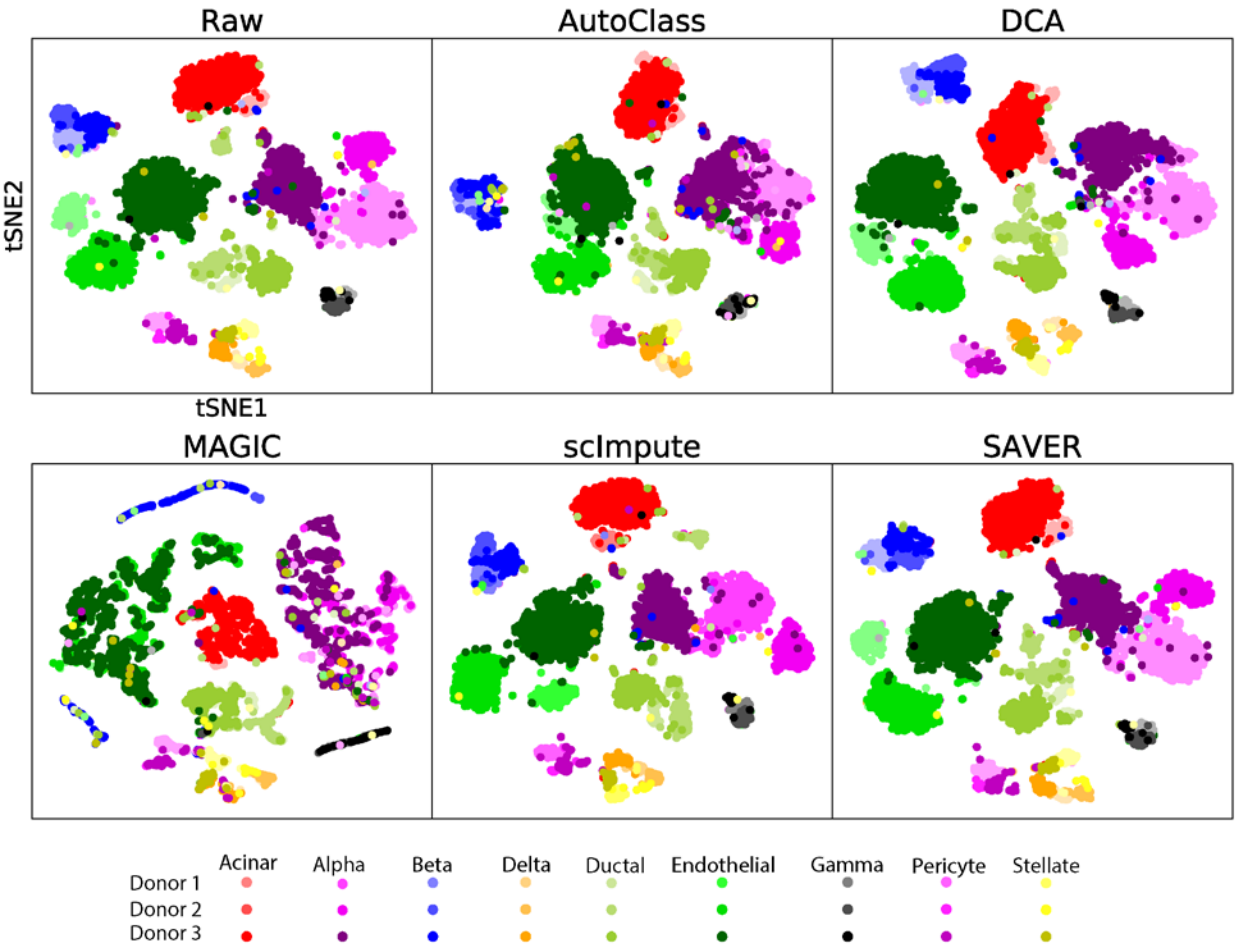


**Supplementary Figure 3** t-SNE plots for raw and imputed data for Baron dataset. Experiments settings are similar to those in Figure 5a.

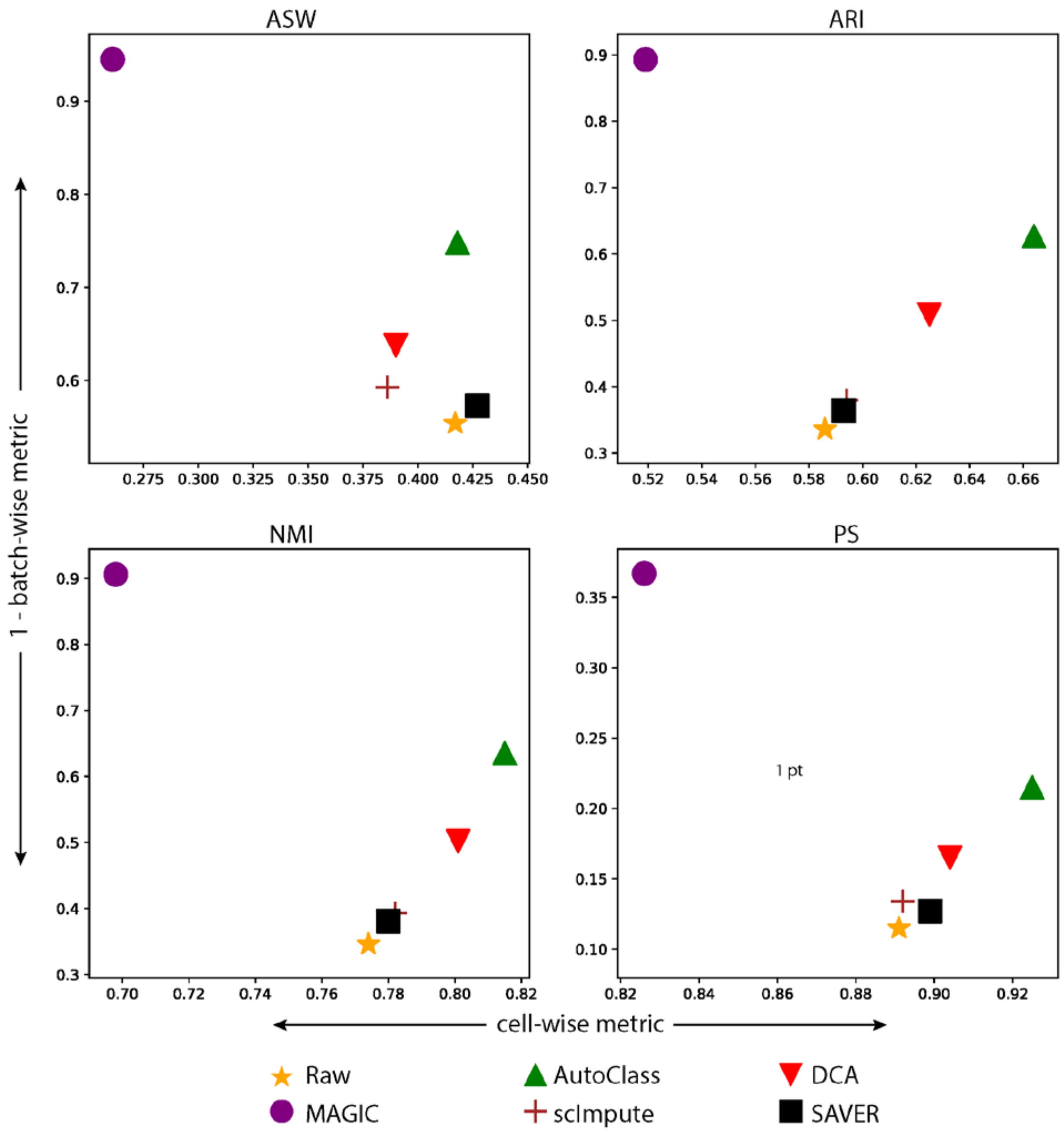


**Supplementary Figure 4** Evaluation of batch and cell type separation in raw and imputed data by four different metrics for Baron dataset. Experiments settings are similar to those in Figure 5b.

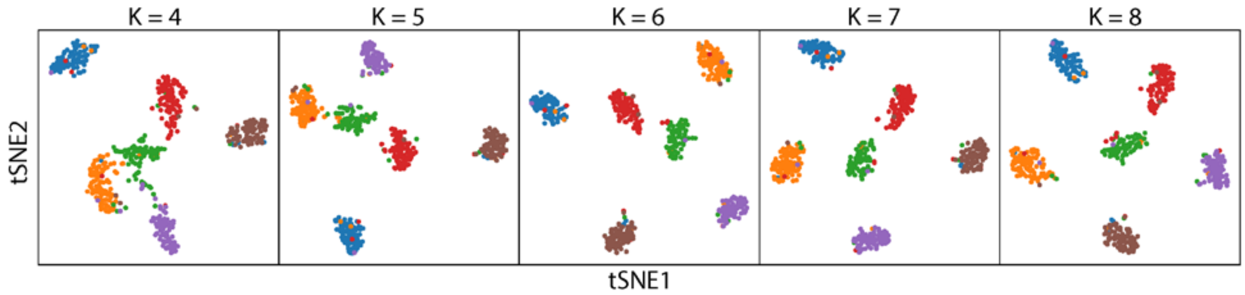


**Supplementary Figure 5** t-SNE plots of Dataset 1 imputed by AutoClass with different $K$ in pre-clustering. Experiments settings are similar to those in Figure 6a.

.

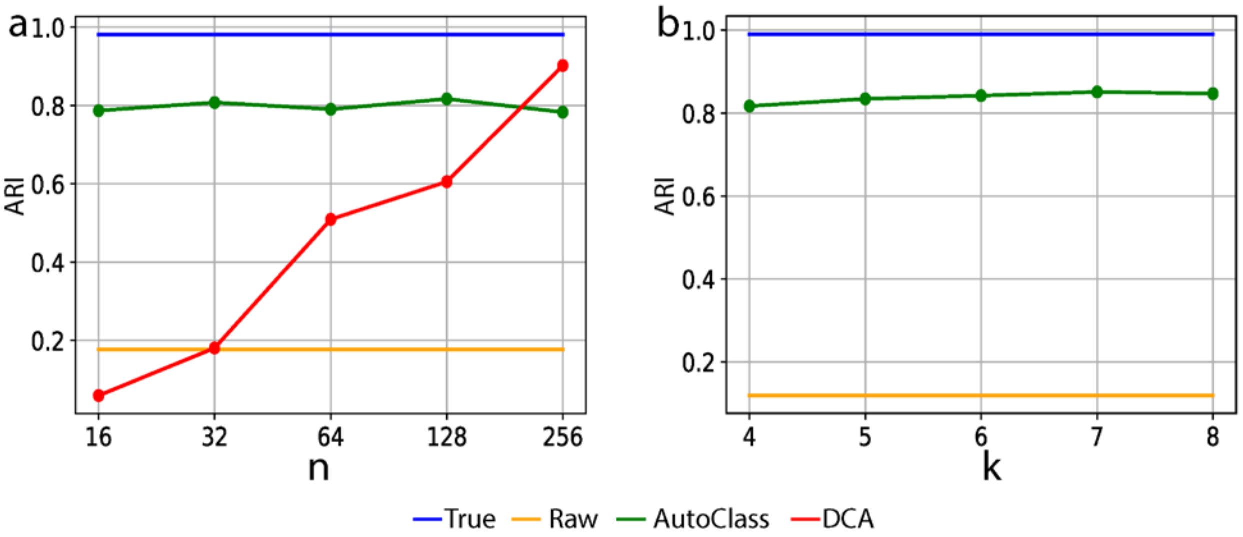


**Supplementary Figure 6** ARI of K-means clustering on t-SNE transformed data for Dataset 1 over **a** different bottleneck layer sizes and **b** different choices of $K$ in pre-clustering of AutoClass (details are described in main text and Methods). Experiments settings are similar to those in Figure 6c and 6d.

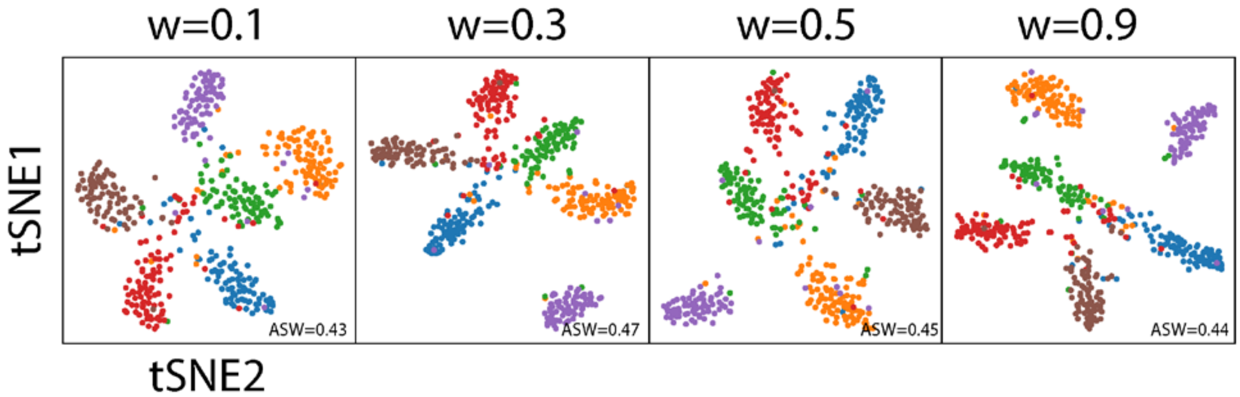


**Supplementary Figure 7** t-SNE plots for Dataset 1 after AutoClass imputation over different classifier weights. Experiments settings are similar to those in Figure 6a.

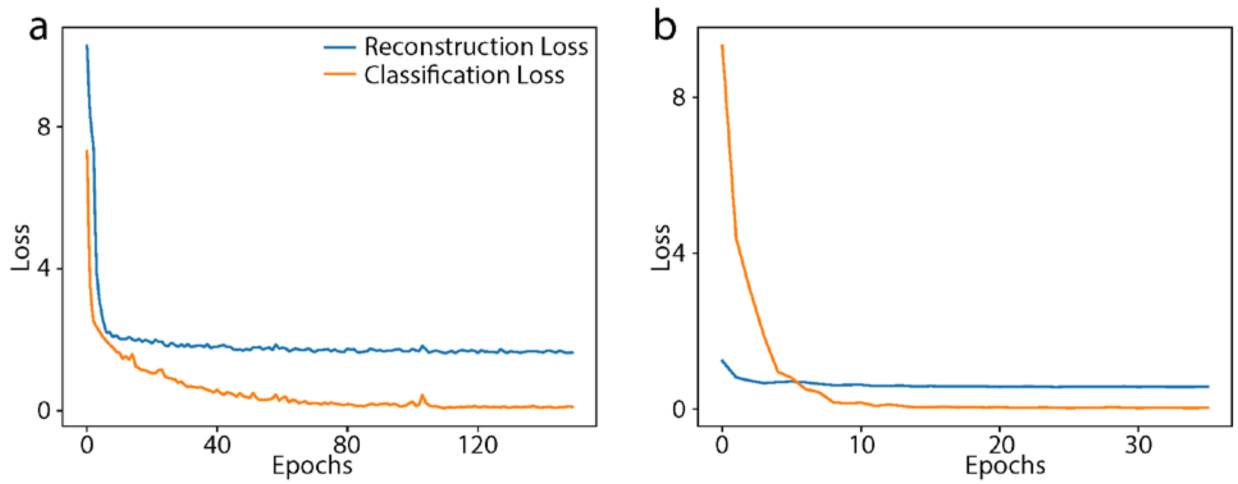


**Supplementary Figure 8** AutoClass loss curves. **a** Dataset 1**. b** Buettner dataset.

| Dataset[Ref] | Description | Cell groups | Sample batches | Cells | % zero counts | AutoClass setting |
|---|---|---|---|---|---|---|
| Baron[16] | Human pancreatic islets cells | 8 | 3 | 7,162 | 82.21 | *dropout_rate=0.3* |
| Villani[24] | Human blood dendritic cells | 4 | 2 | 768 | 47.33 | *dropout_rate=0.3* |
| Lake[18] | Human brain frontal cortex cells | 11 | NA | 8,592 | 73.39 | Default |
| Zeisel[19] | Mouse cortex and hippocampus cells | 9 | NA | 3,005 | 48.58 | Default |
| Buettner[2] | Mouse embryonic stem cells | 3 | NA | 182 | 37.92 | Default |
| Usoskin[17] | Mouse neuronal cells | 4 | NA | 622 | 95.83 | Default |

**Supplementary Table 1** Summary of real scRNA-seq datasets used in this paper.

| Dataset | Genes/Cells | Cell groups | Splatter dropout* | AutoClass setting |
|---|---|---|---|---|
| Dataset 1 | 1000/500 | 6 | -1/5/1.5 | Default |
| Dataset 2 | 1000/500 | 5 | -2.5/2.5 | Default |
| Dataset 8 | 1000/500 | 2 | -1/3 | *num_cluster* = [2,3,4] |

**Supplementary Table 2** Summary for simulated scRNA-seq datasets with dropout noise used in this paper.
*Splatter dropout settings: *dropout.shape/dropout.mid.*

| Dataset | Noise Type | *numpy.random* function | numpy setting | AutoClass setting |
|---|---|---|---|---|
| Dataset 3 | Uniform | *randint* | low=-10, high=10 | Default |
| Dataset 4 | Gaussian | *normal* | loc=0, scale=6 | Default |
| Dataset 5 | Gamma | *gamma* | shape=0.5, scale=12 | Default |
| Dataset 6 | Poisson | *poisson* | lam=6 | Default |
| Dataset 7 | Negative Binomial | *negative_binomial* | n=10, p=0.45 | Default |
| Dataset 9 | Gaussian | *normal* | loc=0, scale=18 | *num_cluster* = [2,3,4] |

**Supplementary Figure 3** Summary for simulated scRNA-seq datasets with non-dropout noises used in this paper. Note Dataset 3-7 were generated from the true data of Dataset 2 above, Dataset 9 from Dataset 8 above.
